# Parkinson’s disease phenotypes in patient specific brain organoids are improved by HP-β-CD treatment

**DOI:** 10.1101/813089

**Authors:** Javier Jarazo, Kyriaki Barmpa, Isabel Rosety, Lisa M. Smits, Jonathan Arias-Fuenzalida, Jonas Walter, Gemma Gomez-Giro, Anna S. Monzel, Xiaobing Qing, Gerald Cruciani, Ibrahim Boussaad, Christian Jäger, Aleksandar Rakovic, Emanuel Berger, Silvia Bolognin, Paul M. A. Antony, Christine Klein, Rejko Krüger, Philip Seibler, Jens C. Schwamborn

**Affiliations:** Developmental and Cellular Biology, Luxembourg Centre for Systems Biomedicine University of Luxembourg, 7 avenue des Hauts-Fourneaux, Esch-sur-Alzette, Luxembourg; Experimental Neurobiology, Luxembourg Centre for Systems Biomedicine University of Luxembourg, 7 avenue des Hauts-Fourneaux, Esch-sur-Alzette, Luxembourg; Metabolomics Platform, Enzymology and Metabolism, Luxembourg Centre for Systems Biomedicine University of Luxembourg, 7 avenue des Hauts-Fourneaux, Esch-sur-Alzette, Luxembourg; Transversal Translational Medicine, Luxembourg Institute of Health (LIH), Luxembourg, Luxembourg; Centre Hospitalier de Luxembourg (CHL), Parkinson Research Clinic, Luxembourg, Luxembourg; Institute of Neurogenetics, University of Lübeck, D-23538, Lübeck, Germany

## Abstract

The etiology of Parkinson’s disease (PD) is only partially understood despite the fact that environmental causes, risk factors, and specific gene mutations are contributors to the disease. Biallelic mutations in the *PTEN-induced putative kinase 1* (*PINK1*) gene involved in mitochondrial homeostasis, vesicle trafficking, and autophagy, are sufficient to cause PD. By comparing PD patient-derived cells, we show differences in their energetic profile, imbalanced proliferation, apoptosis, mitophagy, and a reduced differentiation efficiency to dopaminergic neurons compared to control cells. Using CRISPR/Cas9 gene editing, correction of a patient’s point mutation ameliorated the metabolic properties and neuronal firing rates but without reversing the differentiation phenotype. However, treatment with 2-Hydroxypropyl-β-Cyclodextrin (HP-β-CD) increased the mitophagy capacity of neurons leading to an improved dopaminergic differentiation of patient specific neurons in midbrain organoids. In conclusion, we show that treatment with a repurposed compound is sufficient for restoring dopaminergic differentiation of PD patient-derived cells.

## Introduction

Neurodegenerative diseases pose a great threat to aging populations (Gammon, 2014). Due to the lack of disease-modifying therapies, patients suffering from PD have to rely on symptomatic treatments (Lozano et al, 2018). Affected genes, reported as risk or causative factors of PD, control major cellular processes such as cell proliferation, membrane trafficking, mitochondrial homeostasis, and autophagy (Brás et al, 2015). Among these, PINK1 is involved in regulating mitochondrial function and morphology by quarantining damaged mitochondria before their degradation as well as triggering the process of mitophagy (Ashrafi & Schwarz, 2012). The fact that individuals with biallelic pathogenic variants in *PINK1* develop PD, shows that an altered mitochondrial function, morphology, and degradation are linked to its pathogenesis (Larsen et al, 2018).

One of the hallmarks of PD is the loss of dopaminergic neurons in the *substantia nigra pars compacta*, but other regions have also been reported to be affected (Goedert et al, 2012). Work in the zebrafish model previously suggested that a loss of function of *PINK1* might lead to a developmental reduction in the amount of dopaminergic neurons (Anichtchik et al, 2008). However, implications of *PINK1* and its PD associated mutations, during the transition from neural precursor cells (NPCs) to differentiated neurons have not yet been evaluated in a human cell model. The use of induced pluripotent stem cells (iPSCs) as a source of biological material for modeling neurodegenerative diseases has proven useful for identifying pathogenic key aspects to target (Ghaffari et al, 2018). Another biotechnological tool that facilitated disease modeling is gene editing by CRISPR/Cas9 technique for the generation of isogenic lines (Kim et al, 2014). Correcting *PINK1* mutations in cellular models would allow to analyze the contribution to cellular phenotypes of the actual mutation versus the patients’ genomic background. Techniques for recapitulating disease phenotypes in culture had a recent advancement with the introduction of organoid cultures (Lancaster & Knoblich, 2014; Monzel et al, 2017). These organ-like structures contain different cell types in a spatially organized fashion, recapitulating at least some of the main functions of the respective organ. Importantly, they have been proven valid models for human diseases (Clevers, 2016; Lancaster et al, 2013; Smits et al, 2019).

In this study, PD patients with *PINK1* mutations and healthy individuals’ iPSCs were differentiated into a neuroepithelial stem cell state (NESC), and later into neurons. Different features such as proliferation capacity, apoptosis, and differentiation efficiency were automatically analyzed using computational algorithms for pattern recognition through high-content image analysis. We demonstrated that the differentiation efficiency of patient-derived NESCs is reduced while maintaining an increased proliferative activity upon neuronal differentiation and exhibiting increased apoptosis of dopaminergic neurons. Using extracellular flux analysis and microelectrode array, we assessed the energetic profile of NESCs and the firing activity of differentiated neurons which were altered in patient-derived cells. Moreover, using a pH-sensitive reporter tagging a mitochondrial protein we observed a reduced autophagy and mitophagy capacity. Interestingly, gene correcting the patients’ mutation with the CRISPR/Cas9 system improved the mitochondrial activity and firing rate but not the differentiation efficiency of the isogenic controls. While treatment with the compound HP-β-CD resolved the mitophagy impairment and improved the dopaminergic differentiation in patient-derived cells.

## Results

### Reduced dopaminergic differentiation, increased proliferation and apoptosis in patient-specific cells

To evaluate whether impaired mitochondrial function could affect dopaminergic differentiation, hiPSCs were derived from three patients carrying a mutation in *PINK1*: two carrying p.Q456X (rs45539432) (Seibler et al, 2011) and one carrying p.I368N (rs774647122) and three age- and gender-matched controls (Fig EV1A). Human iPSCs were further differentiated into a stable neural precursor state (neuroepithelial stem cells, NESCs) following a previously published approach (Reinhardt et al, 2013), used as a starting population for studying dopaminergic differentiation efficiency (Fig EV1B). For longitudinal data collection, day 7, 14 and 21 of differentiation were selected as early time points for the quantification of the dopaminergic neuron marker tyrosine hydroxylase (TH). An automated image analysis algorithm was developed to segment and quantify the proportion of neuron-specific class III β-tubulin (TUBB3)-positive signal that colocalizes with TH, and to quantify the nuclear volume as a normalization parameter. Patient-derived NESCs showed a reduced to differentiate into TH-positive (TH+) neurons while maintaining the same level of overall neuronal differentiation compared to controls (Fig 1A and B). Interestingly, the dopaminergic differentiation deficiency worsened at later time points in patient-derived cells (Fig 1B). In parallel, an increased total nuclear area per imaged field in patient cells was also observed (Fig 1B), thus potential proliferation differences were further investigated. To confirm the impaired dopaminergic differentiation in a different culture system, we differentiated NESCs in a 3D microfluidic environment observing the same pattern (Fig EV1C). Assessment of the proliferation marker Ki67 and the apoptotic marker cleaved poly(ADP-ribose) polymerase (cPARP) were performed after 7, 14, and 21 days of differentiation (Fig 1C-F). Patient-derived cells maintained their proliferative capacity after induction of differentiation through a longer period than control cells (Fig 1C and D). This can be related to an impaired exit of the stem cell state while maintaining their proliferative capacity. In addition, patient-derived TH+ neurons showed an increase in cPARP signal (Fig 1E and F). Thus, the impaired differentiation seen in *PINK1* mutant patient-derived cells is caused by a combination of both, stem cells having a lower differentiation efficiency, and TH+ neurons having an increased apoptosis.

**Figure 1.**
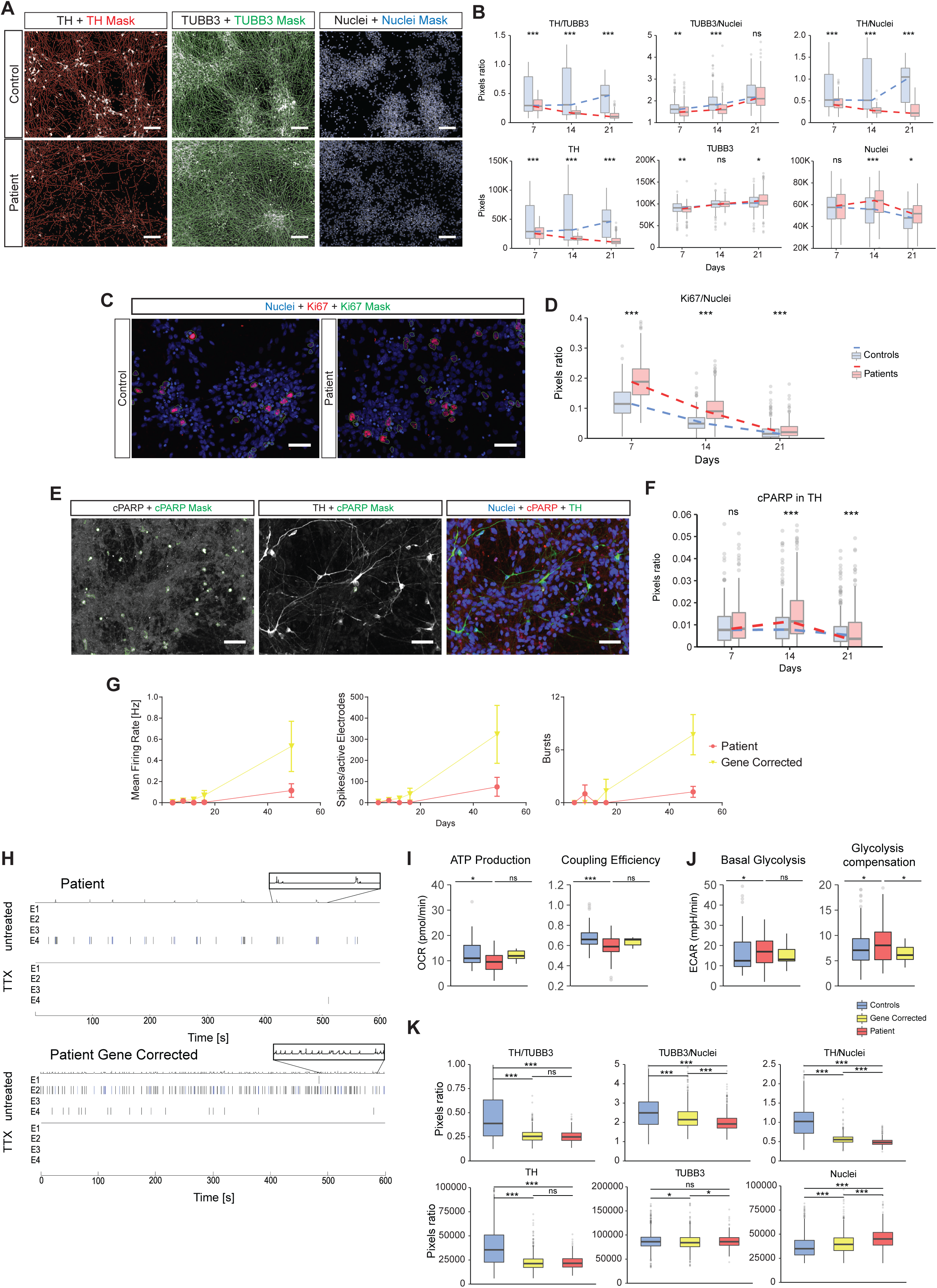
Impaired differentiation of neural stem cells of patient carrying *PINK1* mutations not rescued by gene correction. **A**, Images representing the median values of a 14 day differentiation neuronal culture of controls and patients groups. Raw images of the markers TH, TUBB3 and Hoechst are presented with its respective perimeter mask (scale bar = 100μm). **B**, Quantification of TH, TUBB3 and Hoechst at time points 7, 14 and 21 after the induction of differentiation. Pixel quantification (lower panel) with their respective ratios (upper panel). Acquisition was performed at 20X sampling randomly 15 fields per well. 5 wells of controls and 5 wells of patients were acquired per replicate. *n*= 3 independent replicates using all the lines were performed. Total fields of controls (fc) and of patients (fp) analysed: time point 7 (fc=219; fp=215), 14 (fc=209; fp=219) and 21 (fc=207; fp=220). **C**, Images representing the median values of proliferation marker Ki67 and Hoechst of control and patient-derived cells at day 7 of differentiation (scale bar = 50μm). **D**, Quantification of Ki67 at time points 7, 14 and 21 after the induction of differentiation, normalized to the nuclear area. Acquisition was performed at 20X sampling randomly 15 fields per well. 10 wells of controls and 10 wells of patients were acquired per replicate per time point. *n*= 3 independent replicates using all the lines were performed. Image analysed per time point: 7 (fc=425; fp=421), 14 (fc=411; fp=424), and 21 (fc=416; fp=432). **E**, Representative images of apoptotic marker cleaved PARP (cPARP), dopaminergic marker TH and Hoechst at day 14 of differentiation (scale bar = 50μm). **F**, Quantification of cPARP within TH area at time points 7, 14 and 21 after the induction of differentiation, normalized to TH area. Acquisition was performed at 20X sampling randomly 15 fields per well. 10 wells of controls and 10 wells of patients were acquired per replicate per time point. *n*= 3 independent replicates using all the lines were performed. Image analysed per time point: 7 (fc=219; fp=215), 14 (fc=209; fp=219) and 21 (fc=207; fp=220). **F**, Evaluation of spontaneous neuronal firing by microelectrode measurements (MEA) represented by the mean firing rate, the number of spikes and burst of the neuronal network between patient and gene-corrected cells during differentiation. Data is pooled from three replicates (*n*=3). **H**, Spike raster plots for single electrodes before and after treatment with the voltage-dependent sodium channel blocker tetrodotoxin (TTX). Data is pooled from three replicates (*n*=3). **I**, Extracellular flux analysis (Seahorse) in NESCs for evaluating mitochondrial respiratory capacity and efficiency between controls, patient-derived and gene corrected cells. Data is pooled from three replicates (*n*=3). **J**, Extracellular flux analysis (Seahorse) in NESCs for evaluating glycolytic activity. Data is pooled from three replicates (*n*=3). **K**, Quantification of the markers TH, TUBB3 and Hoechst in a 14 day differentiation neuronal culture with their respective ratios and comparison between patient, gene corrected and control derived neurons. Image analysed: fc=1868, fields gene corrected =796, fp =416. Three independent replicates were performed using all of the lines for all of the panels in this figure, except for panels **G** and **H** where Patient line1 and Patient gene-corrected lines were used. Statistical analysis was performed using a two sample two sided Kolmogorov-Smirnov test for panels **B**, **D**, **F** and **K**. Statistical analysis was performed using Kruskall-Wallis and Dunn’s test for multiple comparisons for panels **I**-**J**. Adjustment of the p-value for multiple tests was performed using Benjamini-Hochberg. \**P*<0.05, \*\**P* <0.01, \*\*\**P* <0.001; ns, not significant.

### Gene correction restored energetic profile but not differentiation efficiency

In order to evaluate the effect of the point mutation in PD patient-derived cells, homozygous correction of the g.20655C>T (p.Gln456Ter) mutation in *PINK1* was performed using FACS-assisted CRIPSR/Cas9 editing (FACE) (Arias-Fuenzalida et al, 2017; Jarazo et al, 2019). Different phenotypic assays were established to assess the effect of the gene editing. Microelectrode array (MEA) measurements demonstrated that the firing activity and the network burst firing in neurons were increased after gene correction (Fig 1G and H, and Fig EV2A). Mitochondrial physiology was metabolically analyzed by extracellular flux analysis (Seahorse) (Fig 1I and J, and Fig EV2B and C). This analysis indicated that the control lines have a higher ATP production from the electron transport chain (ETC) leading to an increased coupling efficiency (Fig 1I). Patient-derived NESCs, in contrast, have a significantly higher glycolytic activity (Fig 1J). These observations are in line with the increased proliferation and the reduced probability of patient-derived NESCs to enter differentiation phases that require a high activity of the ETC. Gene correction of the p.Q456X mutation reduced the compensatory glycolysis response, and improved the metabolic shift and the impaired mitochondrial activity observed in patient cells with a certain tendency to reduce the dependence on glycolysis to generate ATP (Fig 1I and J). Reduction of proliferation upon differentiation was also observed, through a reduction in total nuclei area per imaged field (Fig 1K). Gene correction seems to trigger a metabolic change that allows the switch from stem cells to differentiated cells(Zhang et al, 2012). However, gene correction was not sufficient to increase the proportion of dopaminergic neurons within the neuronal population (Fig 1K).

### *PINK1* patient-specific neurons present a reduced mitophagy capacity

To further understand the mitochondrial dynamics in these cells, we generated a pair of stable patient (*PINK1*, p.I368N) and a control line expressing the Rosella construct bound to microtubule-associated proteins 1A/1B light chain 3B (LC3) or to ATP synthase F1 subunit gamma (ATP5C1), to evaluate the autophagy status and mitochondrial degradation by mitophagy (Arias-Fuenzalida et al, 2019; Sargsyan et al, 2015) (Fig 2). Rosella cells were generated at the hiPSCs state, purified, and further differentiated into NESCs and neurons. Already at the hiPSC level, overall autophagy is reduced in patient-derived cells, shown by a low count of early and late stage autophagy structures (Fig 2A and B). Longitudinal measurements throughout differentiation and automated image analysis revealed significantly fewer mitophagy events in patients’ neurons (Fig 2C and D). However, the mean size of a single mitophagy events was larger (Fig 2D). The observed changes in mitophagy due to p.I368N mutation in *PINK1*, highlight the importance of PINK1 in mitochondrial quality control, especially during the functional and morphological changes occurring during neuronal development. The mitophagy events observed in PINK1 patient-derived cells might be mediated by PINK1-Parkin-independent mitophagy pathways (McWilliams et al, 2018). In order to assess the autophagy pathway as a potential target for small molecules therapy, several known modulators as well as stressors were evaluated as we previously reported (Arias-Fuenzalida et al, 2019). Patient-derived cells treated with rapamycin showed an increase in the frequency of phagophores and autophagic vacuoles, to similar levels as those observed in controls (Fig 2E).

**Figure 2.**
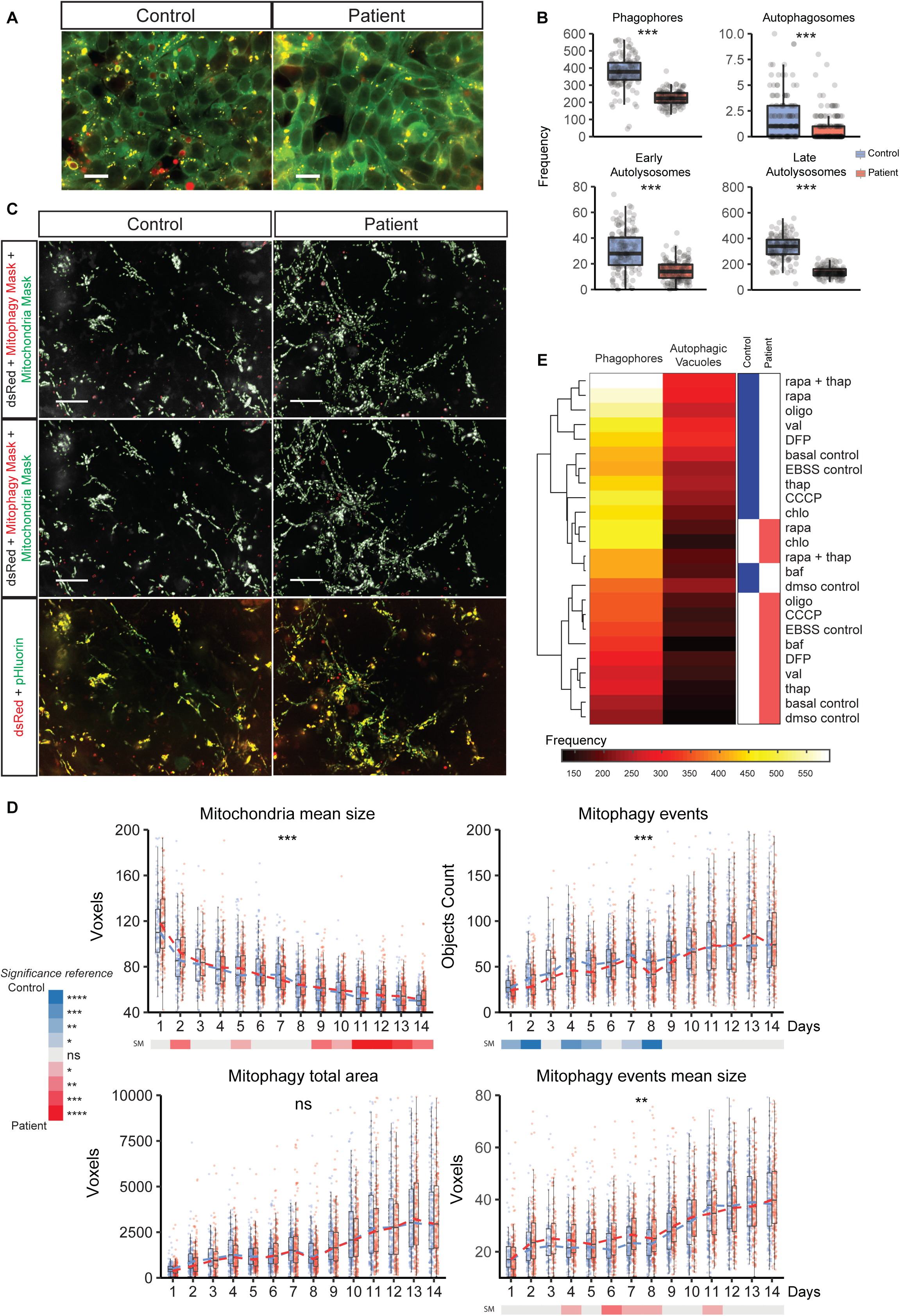
Reduced mitophagy capacity during early neuronal development. **A**, Representative images of hiPSC carrying the Rosella construct targeting LC3 for control and patient-derived cells (scale bar = 20μm). **B**, Absolute quantification of phagophores, autophagosomes, early autolysosomes and late autolysosomes, for controls and patient derived hiPSC. All structures were measured under basal conditions. Acquisition was performed at 60X sampling randomly. *n*= 3 independent replicates were performed. Image analysed: fc=131 and fp=131. **C**, Images representing the median values of neurons tagged with the Rosella construct for depicting mitophagy events at day 8 of differentiation showing the dsRed and pHluorin raw signal (with their corresponding masks) in the left and center panel respectively, of both control and patient-derived cells. A merged image of both channels is shown in the right panel (scale bar = 20μm). **D**, Time series quantification of the mitophagy capacity during neuronal differentiation for 14 days. Different properties of mitochondria and mitophagy events were assessed. Measurements were performed once a day during the entire differentiation protocol. Each dot represents one field fields of controls =97-217 and fields of patients =126-224 range measured per day for 14 days. Acquisition was performed at 60X sampling randomly 15 fields per well. 5 wells of Control 1 and 5 wells of Patient 3 were acquired per replicate (*n*=3, independent replicates). **E**, Heatmap clustering for control and patient derived cells across all mitophagy and autophagy modulating treatments. Scale in absolute event frequency of phagophores or autophagic vacuoles detected. Each dot in the graphs represents one field imaged. Three independent replicates were performed using Control 1 and Patient 3 lines. Statistical analysis for panel **B** was performed using Kruskall-Wallis and Dunn’s test for multiple comparisons. Statistical analysis for panel **D** was performed using a non-parametric test for repeated measures in factorial design (nparLD). Adjustment of the p-value for multiple tests was performed using Benjamini-Hochberg. \**P*<0.05, \*\**P* <0.01, \*\*\**P* <0.001, \*\*\*\**P* <0.0001; ns, not significant.

### Treatment with HP-β-CD increases transcription factor EB (TFEB) nuclear translocation in neurons and dopaminergic differentiation efficiency of brain organoids

Due to the observed altered mitophagy pattern, we explored ways for modulating autophagy with repurposed compounds. It has been reported that treatment with the compound 2-Hydroxypropyl-β-Cyclodextrin (HP-β-CD) modulates autophagy by increasing TFEB nuclear translocation in human neuroglioma cells (Kilpatrick et al, 2015; Martini-Stoica et al, 2016; Song et al, 2014). Treatment with HP-β-CD during the entire differentiation process was able to increase the proportion of nuclei that colocalized with TFEB in a 2D neuronal culture (Fig 3A and B). Patient-derived neurons already presented an elevated level of nuclear TFEB in untreated conditions, suggesting that there is a stronger demand for autophagy proteins (Fig 3B). Moreover, HP-β-CD upregulated the gene expression of FK506 binding protein 8 (FKBP8) and FUN14 domain-containing protein 1 (FUNDC1). Both are outer mitochondrial membrane (OMM)-anchored proteins that present the LC3 interacting region (LIR) motif mediating mitophagy. Interestingly, both genes are down-regulated in patient-derived cells (Fig 3C). Increased nuclear presence of TFEB and expression of the mitophagy receptors FUNDC1 and FKBP8, led to an increase in mitophagy (Fig 3D). In order to evaluate the impact of the treatment in dopaminergic differentiation, midbrain organoids derived according to a published protocol (Monzel et al, 2017), were cultured with different concentrations of HP-β-CD during 30 days of differentiation (Fig 3E). An automated image analysis algorithm was developed to quantify the efficiency of differentiation in 3D structures (Fig EV3A). The proportion of dopaminergic neurons in patient-derived brain organoids was reduced compared to control individuals under basal conditions, while treatment with HP-β-CD increased the proportion of TH+ cells without changing the amount of TUBB3+ neurons (Fig 3E and F). To assess the use of HP-β-CD as a potential general treatment for PD-causing mutations, we extended the treatment to patient-specific neurons carrying the homozygous p.R275W (rs34424986) *Parkin RBR E3 ubiquitin protein ligase* (*PRKN)* mutation. Parkin is known to be a downstream effector of PINK1 induction of mitophagy and alterations can lead to mitochondrial alterations and TH+ neuronal loss (Shaltouki et al, 2015). Also, in these cells, HP-β-CD induced an increase in the amount of dopaminergic neurons, indicating that the potential application of this treatment could be broader (Fig EV3B). We further characterized the composition of the used HP-β-CD mixture and determined that the most frequent isomer is one with six degrees of substitution (DS) (Fig EV4A-C). Knowledge of the HP-β-CD composition will leverage application in future clinical trials and facilitate comparisons of treatment results between different neurodegenerative diseases (Yergey et al, 2017).

**Figure 3.**
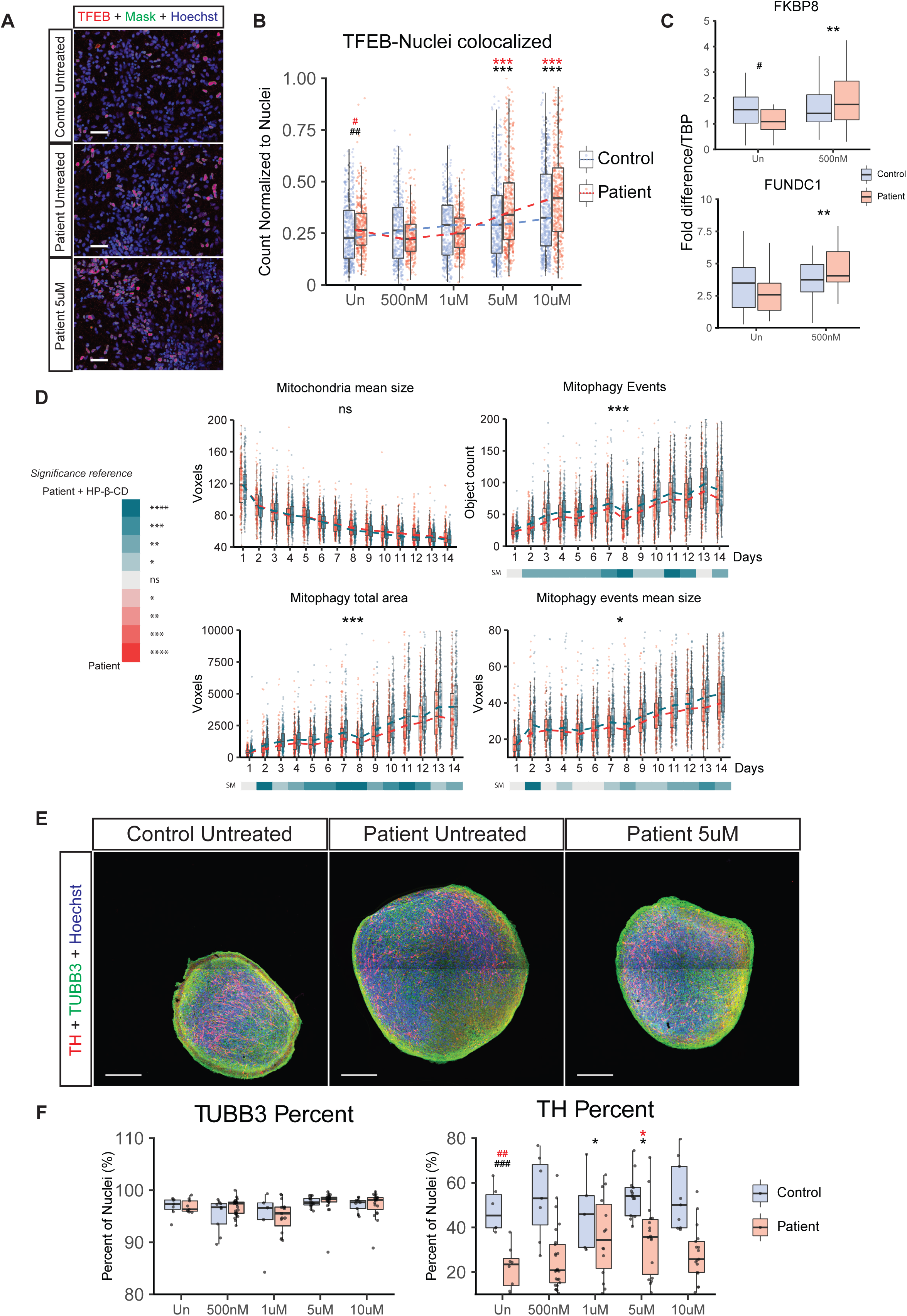
Treatment with HP-β-CD improves neuronal differentiation by increasing autophagy. **A**, Representative images of differentiated neurons stained for TFEB (scale bar = 50μm). **B**, Quantification of the colocalization between TFEB and Nuclei signal with the different treatment concentrations. Each dot represents one field analysed of the 540 fields per category (control or patient) per condition, acquired over *n*=3 independent replicates using all lines. **C**, Relative quantification of 14 day neurons’ gene expression of FKBP8 and FUNDC1 against housekeeping gene (TBP) in treated and untreated conditions over *n*=3 independent replicates. **D**, Time series quantification and comparison of the mitophagy capacity during neuronal differentiation for 14 days between untreated and HP-β-CD treated patient-derived neurons. Different properties of mitochondria and mitophagy events were assessed. Measurements were performed once a day during the entire differentiation protocol. Each dot represents one field analysed over three replicates (*n*=3, fields control -fc- =157-225 and fields patient -fp- =215-225 range measured per day for 14 days. Control 1 and Patient 3 lines were used. **E**, Representative images of control, patient and patient treated derived organoids at 30 days of differentiation (scale bar = 200μm). **F**, Quantification of the markers TH, TUBB3 and Hoechst. Each dot represents one section analysed over *n*=3 replicates. Sections analysed: control (7, 9, 5, 14, and 9) and patient (9, 25, 15, 18, and 19) respectively for the different treatments (Un, 500nM, 1uM, 5uM, and 10uM). Control 1, Patient 1, and Patient 3 lines were used. For panel **F** statistical analysis was performed using a non-parametric test for repeated measures in factorial design (nparLD). For the remainder of the panels, statistical analysis was performed using Kruskall-Wallis and Dunn’s test for multiple comparisons. Adjustment of the p-value for multiple tests was performed using Benjamini-Hochberg (BH). In case that the BH adjustment was performed, the adjusted significance are represented in red. Comparisons between control untreated and patient untreated are presented with #. Comparison between the patient untreated condition, and the different treatment concentrations are represented with *. \**P*<0.05, \*\**P* <0.01, \*\*\**P* <0.001, \*\*\*\**P* <0.0001; ns, not significant. Significance hashtag represent: *P*<0.05 #, *P* <0.01 ##, *P* <0.001 ###; ns stands for not significant. Un, Untreated.

## Discussion

Our findings suggest that a loss-of-function mutation in *PINK1* affects the transition between a neural precursor state and a differentiated dopaminergic neuron (Grünewald et al, 2007) (Fig 1). A previous report showed no difference between a patient line carrying the p.Q456X mutation and control lines, however a different differentiation protocol was used (Chung et al, 2016). Patient-derived cells remained highly proliferative upon induction of differentiation (Fig 1C and D). This matches previous reports showing that the loss of PINK1 activity triggers an increase in glycolysis via a reactive oxygen species (ROS)-mediated stabilization of hypoxia-inducible factor-1α (HIF1α), which has been associated with the Warburg effect (Agnihotri et al, 2016; Requejo-Aguilar et al, 2014). Furthermore, it is in agreement with the finding that PINK1 regulates the metabolic shift away from glycolysis (accompanying manuscript from Bus et al.). Those cells that manage to differentiate into TH+ neurons presented higher levels of apoptosis, showing that the proportion of dopaminergic neurons is affected by a reduced differentiation and an increased cell death in these PD-related mutations (Fig 1E and F). Increased expression of apoptotic and necroptotic markers in the context of PD was previously reported (Iannielli et al, 2018; Michel et al, 2016; Zhang et al, 2017). This can be correlated with an improper metabolic status in mature neurons from PINK1 patients, since oxidative stress mediated neurodegeneration could come from two concomitant origins: ROS from altered mitochondria, and a reduction of NADPH content (and hence a reduced regeneration of glutathione, GSH) due to a compensatory decrease of the pentose-phosphate pathway to increase the glycolytic rate for producing enough ATP (Dias et al, 2013; Herrero-Mendez et al, 2009). Gene correction of *PINK1* allowed neural precursor cells to reduce their dependence on glycolysis, and to increase the firing activity in differentiated neurons (Fig 1G and H). However, lack of improvement in dopaminergic differentiation after gene correction indicates that the patients’ genetic background (Soldner et al, 2011) or acquired mtDNA mutations (Dölle et al, 2016) might play an important role in this phenotype. The susceptibility for developing PD as well as other diseases, has been reported to be different in genetically identical individuals, with epigenetics playing an important role in the onset of the disease (Castillo-Fernandez et al, 2014; Malki et al, 2016; Woodard et al, 2014; Young et al, 2017).

The autophagy pathway is a common process impaired in PD as well as in other neurodegenerative diseases (Guo et al, 2018; Nixon, 2013). PINK1 is directly involved in macroautophagy of mitochondria, known as mitophagy. Impaired mitochondrial turnover, due to a reduced mitophagy activity, was observed in patient cells (Fig 2C and D) in accordance with a previous report (Liu et al, 2011). This altered mitophagy balance occurred simultaneously with an overall impaired autophagy (Fig 2A and B). Using known modulators of autophagy, we modified the quantity of phagophores and autophagic vacuoles in patients’ cells resembling those observed in control cells (Fig 2E). Regulation of autophagy with rapamycin led to a clustering of patient’s together with control’s cells, matching the previously reported effects (Li et al, 2014), and pointing at the autophagy pathway as a candidate to be targeted for compound screening. Rapamycin inhibits the mammalian target of rapamycin (mTOR), which modulates autophagy by reducing TFEB’s nuclear translocation (Martini-Stoica et al, 2016). Nuclear translocation of TFEB regulates the coordinated lysosomal expression and regulation (CLEAR) pathway leading to upregulation of autophagy/lysosomal genes and an increase in autophagy (Martini-Stoica et al, 2016; Sardiello, 2018; Settembre et al, 2013). Due to this close connection, we looked into modulating TFEB activity with HP-β-CD (Song et al, 2014).

Treatment with different concentrations of HP-β-CD increased the presence of TFEB in the nuclei of patient-derived neurons (Fig 3A and B). We observed that HP-β-CD also up-regulated gene expression of FUNDC1 and FKBP8, OMM proteins that recruit the autophagy machinery to degrade altered mitochondria (Fig 3C). FKBP8 also presents anti-apoptotic activity by recruiting Bcl-2 and Bcl-XL to the mitochondria, and also is an inhibitor of mTOR (Bhujabal et al, 2017). In order to assess whether this had a direct impact in mitophagy, patient-derived cells tagged with the ATPC51 Rosella construct were treated with HP-β-CD during the transition between NESCs and 14 day differentiated neurons. The treatment produced a higher number of mitophagy events as well as an increased total area of mitophagy, without modifying the mean size of the mitochondrion (Fig 3D). This implies that the effect of the treatment increased directly the rate of degradation rather than modifying the fission/fusion balance for isolating damaged mitochondria to be removed. We then evaluated the effect of the treatment in patient-derived brain organoids, observing that a concentration of 5μM of HP-β-CD kept stable during the differentiation process improved the impaired dopaminergic differentiation observed in the patient organoids (Fig 3E and F). Strikingly, a beneficial effecto of HP-β-CD on the mitochondrial membrane potential phenotype has been detected in the accompanying manuscript (Bus et al.). Furthermore, it has been reported that increasing autophagy via expression of TFEB, has positive effects in other neurodegenerative disease such as Alzheimer’s, Niemann Pick and Gaucher disease by improving the degradation of protein aggregates (Awad et al, 2015; Willett et al, 2017; Zhang & Zhao, 2015). Moreover, treatment with HP-β-CD is currently in clinical trial for treating Niemann Pick disease (NPD, NCT02534844).

Whether it is a lack of subunit C production for cholesterol processing in NPD type C or an increased overflow of damaged mitochondria due to an altered surveillance by PINK1/Parkin in PD, both can lead to an unbalanced autophagy-lysosomal pathway (Martini-Stoica et al, 2016). This emphasizes the need for identifying commonly altered pathways that would facilitate the transfer of effective compounds between neurodegenerative diseases, since different starting points might face the same roadblock. The present work demonstrates that HP-β-CD ameliorates the dopaminergic neuronal loss phenotype, confirming the therapeutic potential of modulating the autophagy/lysosomal pathway in the context of PD.

## Material and Methods

### Reagents and Resources information

Detailed information about the reagents and resources used in this manuscript are summarized in Table 1.

**Table 1.** Detailed information about the reagents and resources used in the paper.

### Information about cell lines

Healthy control 1 and 2 gave written informed consent at the University of Tübingen. Healthy control 1 and 2 were published in (Reinhardt et al, 2013). Healthy control 3 was provided by Bill Skarnes. We thank the Wellcome Trust Sanger institute, its funders, collaborators and Life Tech limited for supporting us with cell lines. This line is now commercially available in the Human Induced Pluripotent Stem Cells Initiative (HipSci). Patient PINK1 1 and Patient PINK1 2 gave written informed consent at the University of Lübeck. Patient 1 and Patient 2 are sisters of a larger family reported in(Hedrich et al, 2006). Patient PINK1 3 and Patient Parkin samples were obtained from Coriell institute, now hold by the NINDS human cell and data repository. From each donor one clone per iPSC line was derived and used in this study.

### hiPSCs culture, NESCs derivation, neurons differentiation

Fibroblast derived from 3 control individuals and 4 PD patients were reprogrammed to hiPSCs as previously described (Warlich et al, 2011). Human iPSCs were cultured in Matrigel (Corning, 354277) coated plates using Essential 8 (E8) medium (ThermoFisher, A1517001) under 5%CO2 in air mixture. Passages were performed using Accutase (Sigma, A6964) depending on confluency and plated in the same conditions supplemented overnight with 10μM ROCK inhibitor Y-27632 (MerckMilipore, 688000). Human NESCs were generated as described elsewhere (Reinhardt et al, 2013). Briefly, low density iPSCs colonies were treated with dispase (MerckMilipore, SCM133) for isolation and placed in a low addition plate. Cells were cultured in different media for 6 days as described in Figure S1B. Maintenance culture was performed with N2B27 media supplemented with 3μM CHIR-99021 (Axon Medchem, CT 99021), 0.75μM purmorphamine (PMA, Enzo Life Science, ALX-420-045-M005) and 150μM ascorbic acid (Sigma, A4544) as previously described (Monzel et al, 2017). Neuronal differentiation was induced by culturing NESCs in N2B27 supplemented with 10 ng/ml hBDNF (Peprotech, 450-02), 10 ng/ml hGDNF (Peprotech, 450-10), 500 μM dbcAMP (Sigma, D0627), 200 μM ascorbic acid, 1 ng/ml TGF-β3 (Peprotech, 100-36E), and 1 μM PMA (Differentiation media 1) for 6 days. Afterwards, the same media without PMA (Differentiation media 2) was used for the duration of the correspondent experiment. Media changes during differentiation were done every third day. For imaging experiments, cells were seeded in a Cell Carrier Ultra (PerkinElmer, 6055300) 96 well plate. Depending on the assay performed, cell density used was 60000cells/well (Differentiation 14 days) or 30000cells/well (Differentiation staging, proliferation, apoptosis and mitochondria morphology). Experiments were carried out in three replicates (*n*=3). Human iPSCs with a passage number between 15 and 20 after colony picking from the reprogramming plate were used for generating hNESCs. Human NESCs with no more than 10 passages after characterization (at passage 5) were assayed or induced to further differentiate. For organoids generation, 9000 hNESCs per well were seeded in an ultra-low cell adhesion (ULA) 96well plate (Corning). Cells were kept for 7 days in maintenance condition and then subjected for 7 days to Differentiation media 1. Media was afterwards changed to Differentiation media 2, and organoids were kept for 30 days from the start of differentiation. Media changes were performed every 2 days.

### Immunocytochemistry

Fixation was done using 4% PFA in 1xPBS, pH 7.4, for 15min at RT. After 3x 1xPBS washing steps, cells were permeabilized using 0.5% Triton X-100 in 1xPBS for 15 min at RT. Blocking was performed by incubating cells in 5% Normal Goat Serum (ThermoFisher, 10000C), and 0.1% Tween20 in 1xPBS (blocking buffer) for 1 hour at RT. Incubation with the first antibodies was done overnight at 4°C in blocking buffer .Dilution of the antibodies used were: TFEB 1:500, Cleaved PARP 1:400, Ki67 1:200, Sox-1 1:200, TUJ1 (Millipore) 1:1000, TUJ1 (Abcam) 1:500, TH (Santa Cruz) 1:500, Nestin 1:600, Sox-2 1:100, Oct-4 1:400, TRA-1-60 1:100, and TRA-1-81 1:100. Incubation with the secondary antibodies was done after 3x 1xPBS washing steps, for 2h at RT in blocking buffer with 1:1000 Hoechst33342. All secondary antibodies were used at 1:1000 dilution. Cells were washed 3x with 1xPBS, covered with 1xPBS and imaged directly after.

### Immunohistochemistry

Processing of organoids was performed as described in (Monzel et al, 2017) with some modifications. Briefly, organoids were fixed with 4 % paraformaldehyde overnight at room temperature and washed 3 times with PBS. Afterwards, they were embedded in 3 % low-melting point agarose (Biozym) in PBS and incubated for 15 min at 37 °C, followed by 30 min incubation at RT. The solid agarose block was covered with PBS and kept overnight at 4 °C. Sections were performed using a vibratome (Leica VT1000s) obtaining 50 µm sections at a speed of 6 and a frequency of 8. Sections were transferred to a 24well plate to perform the staining as free floating sections in 250ul of staining solution per well. Sections were incubated in 2% normal goat serum, 2% bovine serum albumin and 0.5 % Triton X-100 in PBS for 90 minutes on a shaker, and afterwards kept for 48h in a similar solution but with 0.1% Triton X-100 on a shaker for at 4 °C with primary antibodies. Primary antibodies dilutions for the organoids were the following: TH 1:1000 (Abcam), and TUJ1 1:1000 (BioLegend). After incubation, sections were washed 3 times with 500ul PBS. Incubation with the secondary antibody was perform in 2% normal goat serum, 2% bovine serum albumin and 0.05 % Tween-20 in PBS for 2 hours at RT with 1:1000 Hoechst33342. All secondary antibodies were used at 1:1000 dilution. Sections were mounted on Teflon coated slides (De Beer Medicals) with mounting medium (SouthernBiotech) and covered with a 24×50mm cover glass (VWR). Slides were imaged after 24 hours.

### Extracellular flux analysis (SeaHorse measurements)

Human NESCs were seeded in a Matrigel coated XF 96-well plate (Agilent, 102416-100) at a density of 65000 cells per well. Cells were incubated in a normal incubator for 6h for attachment. Media was removed and washed 2x using Assay medium consisting of 1mM Pyruvate (ThermoFisher, 11360-039), 21.25 mM D+glucose (Sigma, G8270), 2mM glutamax (ThermoFisher, 35050061) in DMEM (Sigma, D5030), at 37°C, pH 7.4. For equilibrating the plate, cells were left in assay medium at 37°C in air for 1 hour before running the assay in a Seahorse XFe96 Analyser according to manufacturer instructions. Concentrations of compounds after the injection were 1μM oligomycin (Sigma, 75351), 1μM FCCP (Sigma, C2920), 1μM antimycin A (Sigma, A8674) and 1μM rotenone (Sigma, R8875). Three baseline measures and three measurements after each compound injection were performed. For normalization, DNA was quantified using CyQUANT kit (ThermoFisher, C7026) and absorbance was measured in a Cytation 5 plate reader (BioTek) at 480/52.

### MEA measurements

The Maestro microelectrode array (MEA, Axion BioSystems) system was used to measure spontaneous activity of dopaminergic neurons. 96-well MEA plates were pre-coated with 0.1 mg/ml poly-D-lysine hydrobromide (Sigma, P7886) in 0.1 M borate buffer (Sigma, T9525) for 1h at RT. 1*10^5^ hNESCs were placed onto each array in a Matrigel droplet and incubated for 1h at 37 °C before adding culturing media. Cells were recorded during the course of dopaminergic differentiation at a sampling rate of 12.5 kHz for 10 min at 37 °C. Axion Integrated Studio (AxIS 2.1) was used to process the raw data as previously described (Monzel et al, 2017). Using Axion Integrated Studio (AxIS 2.1), a Butterworth band pass filter with 200-3000 Hz cut-off frequency and a threshold of 6 x SD were set to minimize both false-positives and missed detections. The Neural Metric Tool (Axion BioSystems) was used to analyse the spike raster plots. Electrodes with an average of ≥5 spikes/min were defined as active. The spike count files generated from the recordings were used to calculate the number of spikes/active electrode/measurement. For the pharmacological treatment (*n*=3) neurons were treated with Tetrodotoxin (TTX, Cayman Chemical, 14964, 1 µM) for blocking and hence verifying the neuronal activity. The spike count files generated from the recordings were used to calculate the number of spikes/active electrode.

### Rosella mitophagy reporter

The pH sensor fluorescent protein pHluorin was fused to DsRed and the entire open reading frame of ATP5C1 or LC3 as described in (Arias-Fuenzalida et al, 2019; Sargsyan et al, 2015). The entire cassette was introduced into the AAVS1 safe-harbour locus as previously described using the targeting donor (Addgene 22075) and TALE nucleases (Addgene 35431 and 35432) as described in(Hockemeyer et al, 2009). Human iPSCs were electroporated with a Lonza 4D nucleofector system subunit X (Lonza, AAF-1002X) according to manufacturer instructions, using a P3 primary cells kit (Lonza, V4XP-3024) and pulse CB-150. Compound treatment for modulating autophagy response was performed as described in (Arias-Fuenzalida et al, 2019).

### Gene editing

Gene correction of patient’s point mutation was performed as previously described(Arias-Fuenzalida et al, 2017; Jarazo et al, 2019). Briefly, for designing the gRNAs targeting sequence for PINK1e7 456 candidates were selected in silico (https://portals.broadinstitute.org/gpp/public/analysis-tools/sgrna-design) and inserted into pX330 vector (Addgene 42230) as described in(Ran et al, 2013). Donor constructs were assembled by introducing the correspondent homology arms into donor scaffold containing a positive selection module (PSM) and either EGFP or dTomato (PSM-EGFP and PSM-dTomato) and a blue florescence in the backbone of the plasmid for detecting random integrations (tagBFP), using Gibson assembly. After sorting of yellow colonies in BD FACS ARIA III, removal of the positive selection module (PSM) was performed by transfecting transposase piggyBac excision only mRNA with Stemfect RNA transfection kit (Stemgent, 00-0069) following manufacturer’s protocol. Cell sorting was performed again for isolating cells with a proper removal of the PSM. The gene correction was done in PINK1 line 1 (Supplementary Fig. 1).

### RNA isolation, RT-PCR and qPCR

Total RNA was isolated from cells using miRNeasy® Mini Kit (Qiagen) and treated with RNase-Free DNase Set (Qiagen). cDNA was reverse transcribed using the High-Capacity RNA-to-cDNA™ Kit (Invitrogen). Amplification of the genes of interest by PCR was performed using FKBP8 Iso2 and FunDC1 primers. Amplification of the housekeeping genes by PCR was performed using TBP (TATA-binding protein) primers. SYBR green qPCR was performed using LightCycler 480 SYBR Green I Master (Roche) with each biological sample ran in triplicate. Quantification of gene expression was performed using the LightCycler® 480 Probes software (Roche).

### Compound treatment

Evaluation of the compound treatment was performed using Cell Carrier Ultra plates seeding NESCs at a density of 30000 cells/well. 2-Hydroxypropyl-β-Cyclodextrin (HP-β-CD, Sigma, H-107) was added on every media change at the different concentrations tested and kept throughout the entire differentiation process. Treatment of the organoids was performed in the same way but in a 96well ULA plate format for 30 days of differentiation.

### Image acquisition

Cell carrier Ultra plates were imaged in an automated manner using an OPERA QEHS spinning disk microscope (PerkinElmer). Depending on the experiment images were acquired with a 10x air objective (Differentiation efficiency) or 20x water immersion objective (for apoptosis and proliferation) or a 63x water immersion objective (mitochondria morphology and mitophagy). Imaging of live cells (mitophagy assessment) was performed under normal incubation conditions (37°C, 80% humidity and 5% CO2 in air). Both fluorescence (pHluorin and DsRed) were acquired at the same time using two cameras using bandpass filters (520/35 and 600/40 respectively). For 3D evaluation Z stacks were performed with an interval of 3.2 μm (acquisition of the entire height of the microfluidic chip) or 400nm for the images acquired with the 63x objective. Imaging of the organoids’ sections was performed in the Yokogawa CV8000 using the Search First modality at 4x to identify the position of the sections, and re-scan at 20x air objective for acquiring the final images.

### Image analysis

The image analysis was performed using MatLab, The MatWorks Inc. as previously described (Arias-Fuenzalida et al, 2019; Bolognin et al, 2018).

### RNA isolation, RT-PCR and qPCR

Total RNA was isolated from cells using High Pure RNA isolation kit (Roche). cDNA was reverse transcribed using the Transcriptor High Fidelity cDNA Synthesis Kit (Roche). Amplification of the genes of interest by PCR was performed using FKBP8 Iso2 and FunDC1 primers as shown in STAR Methods. Amplification of the housekeeping genes by PCR was performed using ACTB (b-actin) and TBP (TATA-binding protein) primers. SYBR green qPCR was performed using LightCycler 480 SYBR Green I Master (Roche) with each biological sample running in triplicate. Quantification of gene expression was performed using the LightCycler® 480 Probes software (Roche). More information about qPCR primers and primers are available in supplemental tables.

### Microfluidics culture

Neuroepithelial stem cells were seeded in an OrganoPlate (Mimetas, 9603-400-B) as explained elsewhere (Moreno et al, 2015). Briefly, a cell suspension in liquid Matrigel was prepared at a concentration of 20000 cells/μl of Matrigel. Cells were seeded with a repeating pipette with 1 μl of the solution per chip. The media perfusion was achieved by gravity, with an average fluid flow of 1.5μl/h (Trietsch et al, 2013). For immunocytochemistry, cells were fixed with 4% PFA in 1xPBS, pH 7.4 overnight at 4°C and both antibodies incubation were kept overnight at 4°C.

### Mass Spectrometry

Ion mobility - MS experiments were performed using an Agilent 6560 Ion Mobility – QTOF MS equipped with an Dual Agilent Jet Stream ESI source (Agilent Technologies). The following method is based on a protocol from (Yergey et al, 2017). First, three stock solutions of 2-Hydroxpropyl-β-cyclodextrin (*c* = 1 mg/mL, Sigma, H107) were prepared in 1mol/L ammonium hydroxide solution. Then, samples were diluted in 50:50 (v/v) ACN:H2O+0.1% formic acid to a final concentration of 10 µg/mL. Samples were directly infused into the ion source (positive ion mode) for 2 min using a flow rate of 200 µL/min. Nitrogen was used as drying gas at a temperature of 300 °C, a drying gas flow of 5 L/min, a sheath gas temperature of 350 °C, and a sheath gas flow rate of 11 L/min. The nebulizer gas pressure was set to 35 psig, the MS capillary voltage was 3.5 kV, the nozzle voltage was 1 kV, and the fragmentor was set to 400 V. Data was acquired in a mass range from *m/z* 100 to 3200. The Instrument was operated in IM-QTOF mode and tuned in high resolution mode (slicer position: 5) and Extended Dynamic Range (2GHz). Ions were trapped for 20,000 µs and released every 60 ms with a trap release time of 150 µs. The drift tube was operated with an absolute entrance voltage of 1700 V and an exit voltage of 250 V with a drift tube pressure of 3.94 Torr and a temperature of 31 °C using nitrogen 6.0 as the collision gas. The acquisition settings were adjusted to yield 32 ion mobility transients/frame corresponding to 0.5 frames/sec. External mass calibration as well as SingleFieldCalibration was performed before measurement of each set of samples and according to the manufacture instructions. All data were acquired with Agilent Mass Hunter LC/MS Data Acquisition (ver B.08.00) and analysed with Agilent Mass Hunter IM-MS Browser (ver B.08.00), where all acquisition frames were extracted from a total time frame of 1.8 min (selected time range: 0.1 to 1.9min).

### Statistical Analysis and graphical representation

Statistical analysis performed on each assay is mentioned on each figure legend. All the statistical analysis were performed in R. When residuals of the data were not normally distributed after QQplot evaluation, non-parametric analysis were performed using Kruskal-Wallis (KW) test. If the assumption of identical distributions (same shape) of non-normally distributed was not fulfilled for performing a KW test, a two-sample Kolmogorov-Smirnov (KS) test was used to evaluate if the samples were drawn from significantly different distributed populations between control and patient lines. For the repeated measures studies done with the lines tagged with the Rosella construct, a non-parametric test for repeated measures in factorial design was performed using the R package nparLD with an F1.LD.F1 design (Noguchi et al, 2012). Adjustment of the p-value for multiple tests was performed using Benjamini-Hochberg (BH). In case that the BH adjustment changed the p-value and the decision, the adjusted significance are represented in red. Horizontal lines in dot plots, violin plots and box plots represent the median. Vertical lines in dot plots, violin plots and line plot, and hinges of the box plots represent the first and third quantile (the 25th and 75th percentiles). Whiskers of the box plots extend to 1.5*Inter-quantile range from the hinges. A dot in a dot plot represents a field of a well, except for the microfluidic experiment where each dot represents an entire chip. Significance asterisks represent: *P*<0.05 *, *P* <0.01 **, *P* <0.001 ***; ns stands for not significant. Significance hashtag represent: *P*<0.05 #, *P* <0.01 ##, *P* <0.001 ###; ns stands for not significant.

## Acknowledgments

We thank Prof. Dr. Hans R. Schöler of the Max Planck Institute, Dr. Jared Sterneckert of the CRTD, Prof. Dr. Thomas Gasser of the Hertie Institute in Tübingen, William Skarnes of the Jackson Laboratory and the Coriell Institute for providing cell lines. Prof. T. Graham and A. Sargsyan from the University of Utah for kindly providing us with the Rosella construct. Acquisition of flow cytometry data was supported by the flow cytometry core of the LCSB bio-imaging platform. Zdenka Hodak of the LCSB Metabolomics Platform for providing technical and analytical support. We thank the Disease Modeling Screening Platform from LCSB and LIH for their help with performing automated and high-throughput procedures. This project was funded by the Fonds National de la Recherche (FNR) Luxembourg (CORE, C13/BM/5791363). This is an EU Joint Program - Neurodegenerative Disease Research (JPND) project (INTER/JPND/14/02; INTER/JPND/15/11092422). This project is also supported by the European Union’s Horizon 2020 research and innovation programme under grant agreement No 668738, SysMedPD. The automated screening platform was supported by a PEARL grant of the Luxembourg National Research Fund (FNR) to R.K. J.J. is supported by a Pelican award from the Fondation du Pelican de Mie et Pierre Hippert-Faber. J.J., L.M.S., A.S.M., J.W. and X.Q. were supported by FNR Aides à la Formation-Recherche (AFR). G.GG. was funded by the NCL-Stiftung (Hamburg, Germany). Finally, we also thank the private donors who support our work at the Luxembourg Centre for Systems Biomedicine. A.R., C.K., and P.S. are supported by the DFG (FOR2488).

## Author contributions

J.J. and J.C.S. designed the study. J.J., I.R., L.M.S, J.A.F., J.W., G.G.G., A.S.M., X.Q., G.C., I.B., C.J., P.S., A.R., E.B., S.B. and P.M.A.A. established methodology. J.J., K.B., I.R., L.M.S., J.A.F., J.W., G.G.G., A.S.M., X.Q., G.C., C.J., P.S., A.R., S.B. and P.M.A.A. performed experiments. C.K., R.K. and P.S. supervised. J.J. made the final figures and wrote the manuscript. J.C.S. conceived and supervised the study.

## The paper explained

### Problem

Parkinson’s disease is one of the most prevalent neurodegenerative diseases. However, after more than 150 years since the disease was described, no medical treatment that reduces the loss of dopaminergic neurons is available. One of the reasons for this failure, is that drug candidates were screened in models that do not recapitulate entirely the pathogeny of the disease observed in humans.

### Results

Disease modelling in a dish can help to identify potential targets for developing pharmaceutical therapies to be further tested in vivo. Using a 3D model such as organoids recapitulates the architecture and function of the affected organ in a more reliable way. Here we applied such tools to model Parkinson’s disease. Using patient derived samples we observed that the amount of dopaminergic neurons was reduced upon neuronal differentiation. The impaired dopaminergic differentiation occurred concomitant with a reduced mitophagic capacity that can be linked to the loss of function of PINK1. Gene correction of the point mutation was not able to fully restore these phenotypes to the healthy levels. However, treatment with a repurposed compound (HP-β-CD) was able to improve the differentiation phenotypes observed in vitro.

### Impact

HP-β-CD may prevent the development or progression of the disease through the induction of autophagy.

## Expanded View Figure Legend

**EV Figure 1**

**A**, Table summarizing the lines used in the article. **B**, Time line and procedure for the generation of the lines and quality controls performed in the lines used. hiPSCs characterization scale bar = 100μm. hNESCs characterization scale bar = 500μm. Differentiation evaluation 2D scale bar = 100μm. Differentiation evaluation 3D scale bar = 200μm. **C**, Representative images of a 14 day differentiation neuronal culture on a chip of an OrganoPlate stained for TH, TUBB3 and a nuclear marker (scale bar = 100μm). **D**, Quantification of TH normalized to neuronal area. Each dot represents a chip in an OrganoPlate over *n*=3 replicates (chips control = 84, chips patient = 82). Statistical analysis was performed using a two sample two sided Kolmogorov-Smirnov test. *P<0.05, **P <0.01, ***P <0.001.

**EV Figure 2**

**A**, Representative raw data trace before and after addition of the voltage-dependent sodium channel blocker tetrodotoxin (TTX) showing that the traces observed were produced by firing neurons. **B**, Representative scheme of the mitochondrial stress test profile for mitochondrial respiration and the areas used for the calculations obtained from the extracellular flux analysis. **C**, Representative oxygen consumption rates during the mitochondrial stress test.

**EV Figure 3**

**A**, Representative images of organoid sections with the respective masks identifying TH, TUBB3 and Hoechst. Scale bar = 200μm. **B**, Quantification of the markers TH, TUBB3 and Hoechst in a 14 day differentiation neuronal culture with their respective ratios and comparison between untreated and HP-β-CD treated *PRKN* patient-derived neurons.

**EV Figure 4**

**A**, Ion mobility abundance map (drift time vs. mass-to-charge) of an HP-β-CD sample indicating singly charged “Monomers” (DS_n_) and doubly charged “Homo- and Heterodimer” (2DS_n_ and DS_n_+DS_n-1_) regions. **B**, Extracted mass spectrum of region “Monomers”, showing HP-β-CD monomers with different degrees of substitution (DS_n_, ammonium adduct [M+NH4]^+^). Asterisk indicates the less abundant corresponding protonated molecule [M+H]^+^. **C**, Relative fractional intensity as function of the degree of substitution (DS_n_) in the HP-β-CD sample.

